# SNPnexus COVID: Facilitating the analysis of COVID-19 host genetics

**DOI:** 10.1101/2020.12.18.423439

**Authors:** Jorge Oscanoa, Lavanya Sivapalan, Maryam Abdollahyan, Emanuela Gadaleta, Claude Chelala

## Abstract

The severe acute respiratory syndrome coronavirus 2 (SARS-CoV-2) pandemic has demanded an unprecedented scientific response, with researchers collaborating on a global scale to better understand how host genetics can influence susceptibility to coronavirus infection and the severity of COVID-19 symptoms. The number of projects directed towards sequencing patients’ genomes has increased rapidly during this time with the rate of data generation outpacing the resources available for analysis and biological interpretation of these datasets. SNPnexus COVID is a cutting-edge web-based analytical platform that allows researchers to analyse and interpret the functional implications of genetic variants in COVID-19 patient genomes and to prioritise those that demonstrate clinical utility for the prevention, management and/or treatment of COVID-19. Our resource links to diverse multifactorial datasets and information resources that would require substantial time and computational power to otherwise mine independently. This streamlines biological data interpretation and allows researchers to better understand the multidimensional characteristics of their data. Importantly, SNPnexus COVID is powered by the SNPnexus software and follows its intuitive infrastructure, which precludes the need for programmatic experience in its users.

SNPnexus COVID is freely available at https://www.snp-nexus.org/v4/covid/

## INTRODUCTION

The pandemic caused by the severe acute respiratory syndrome coronavirus 2 (SARS-CoV-2) has presented a crisis for global healthcare systems (1,2). Human SARS-CoV-2 infection can result in coronavirus disease 2019 (COVID-19), which has been characterised as an acute respiratory illness, with most patients displaying flu-like symptoms, such as a fever, cough and dyspnoea (1–3). However, the range and severity of individual symptoms experienced by patients can vary significantly, indicating a role for host genetics in impacting the susceptibility and severity of COVID-19 disease. Whilst most symptomatic infections are known to manifest in mild to moderate respiratory symptoms, severe pneumonia and complications including cytokine release syndrome, which can lead to multi-organ dysfunction, have also been observed in cases worldwide (4,5).

Global initiatives to sequence the genomes of patients with COVID-19 have driven an expanding new field of host genomics research to characterise the genetic determinants of COVID-19 disease (1). The functional annotation and analysis of incoming genomic data, within a clinically relevant turnaround time, is therefore imperative given the importance and urgency of research efforts to understand the biology of SARS-CoV-2 infection and disease.

To address these requirements, we developed SNPnexus COVID. This is a web-based variant annotation tool, powered by the SNPnexus software (6).

## MATERIAL AND METHODS

### Architecture

SNPnexus COVID adopted the SNPnexus software and 3-tiered framework as its underlying query architecture (as described previously (6)). In this framework, the core annotation process is decoupled from the scheduler and the web interface to streamline the analytical workflow, increase query response times and reduce computational burdens, thereby allowing for the management of significantly larger input uploads.

The range of annotations available from SNPnexus COVID is expansive and tailored for the study and interpretation of COVID-19 host genetics. One of the major enhancements to the SNPnexus framework is the ability to handle both single and multiple samples from a sequencing project. This new feature in conjunction with the filtering system and the new annotations, provides a compelling tool for prioritisation of relevant variations potentially involved in the different symptomatologies of COVID-19.

### Data queries

A significant proportion of COVID-19 research has applied whole genome and exome sequencing approaches for host genotyping. Currently, SNPnexus COVID allows users to submit a total of 200,000 variants per single file and up to a maximum of 12 variant files aligned to the hg38 human genome assembly. Alternatively, unfiltered vcf files can be uploaded to SNPnexus COVID provided that the user selects the pre-filtering option to restrict input variants based on gene panels reported to be involved in SARS-CoV-2 virus-host interaction mechanisms from GENCODE and Genomics England (https://www.gencodegenes.org/human/covid19_genes.html, https://panelapp.genomicsengland.co.uk/panels/111/). The annotation of human protein-coding genes is being updated as part of the GENCODE project (7) to include new transcript models for the study of disease-associated variants in patients with severe COVID-19. Similarly, Genomics England PanelApp (8) provides a list of genes associated with susceptibility to viral infection, including SARS-CoV-2.

### Data annotations

SNPnexus COVID collates and integrates information from a broad range of annotation datasets and multifactorial resources (Supplementary Table 1). To the best of our knowledge, there is no COVID-dedicated resource available in the public domain that provides the breadth of data and functionalities offered by SNPnexus COVID.

These user-directed filtering options will be used to filter uploaded vcf files according to the chromosomal loci of target genes for functional annotation. Similar to the existing SNPnexus architecture, users can download queried results in either a per-variant or per-annotation view and/or explore the results online via a range of interactive graphics and tables.

By enabling users to leverage the expansive datasets that are integrated within our tool, through a simple online filtering and query system, SNPnexus COVID offers significant scope for the acceleration of drug discovery efforts.

### Annotation fields for functional characterisation of variants associated with COVID-19 disease

#### Gender-based population frequencies

COVID-19 has been shown to have a higher mortality rate amongst men, as well as in the presence of comorbidities, demonstrating the value of genomic profiling to identify variants that may underpin a possible gender-dependent susceptibility (9,10). Alternate allele frequencies for variants within COVID-19 genes can be queried against existing normal population databases; 1000 Genomes, gnomAD exomes / genomes and HapMap. In SNPnexus COVID, gnomAD annotations allow for gender-based stratification of population variant allele frequencies across different populations.

#### Gene expression queries

Evaluating the impact of genetic variation on the expression of genes involved in SARS-CoV-2-human interaction is integral for the identification of functional coding variants, which may play a role in the variability of SARS-CoV-2 transmission between populations and/or the severity of COVID-19 symptoms (11). GTEx tissue-specific and global gene expression profiles for candidate COVID-19 genes are available in SNPnexus COVID (GTEx v8). Moreover, users can analyse expression quantitative trait loci (eQTL) affecting target gene expression across different cell types and tissues, to identify genetic determinants of expression.

#### Network characterisation of Protein-Protein and Drug-Target interactions

The characterisation of drug-target interaction networks can provide an important tool to identify potential targets amenable to treatment with existing drugs. The networks available from SNPnexus COVID, based on the protein-protein and protein-drug interactions, will help users gain a better understanding of the host interactome (12,13). Variants within candidate COVID-19 genes can be queried against the DrugBank database, for the analysis of potential genotype-driven therapeutic targets.

#### Pathway analysis

Data interpretation in context of biological systems is vital to understand biological features defining a phenotype. Pathway analyses are available to further help understand the host tissue response to COVID-19. Variants in the query set are mapped to their corresponding genes, which, in turn, are linked to biological pathways in the Reactome Pathway Database. An interactive table and Voronoi diagram provide users with a landscape of pathways significantly enriched in the uploaded variant set.

### Documentation

A description of input/output formats are available from the online User Guide, with practical demonstrations of single and multi-sample cohort queries also provided as videos. Results from the practical demonstrations are available to query from https://www.snp-nexus.org/v4/covid/results/sarscov2/ and https://www.snp-nexus.org/v4/covid/cohort/covid_cohort/.

## RESULTS

### Case of Use

Innovative tools and methods are essential to decode the sequencing data and identify actionable opportunities for intervention aimed at optimising patient outcomes and their safety. To test the performance of SNPnexus COVID and its ability to analyse and prioritise actionable targets we conducted a query using variant files obtained from The COVID-19 Host Genetics Initiative (https://www.covid19hg.org/).

These files comprise filtered variants aligned to the hg38 genome assembly from the B1, C1 and C2 cohorts (https://www.covid19hg.org/results/) and represent hospitalised/non-hospitalised COVID-19 infected patients as well a COVID-19 negative population.

A summarised view of variant features allows users to inspect variant classification and type, SNV class, variant distributions across cohorts/patients and top mutated genes (Figure 1A). Mutations in *FYCO1*, *IFNAR2* and *TYK2*, genes linked to increased susceptibility to severe viral infections, including SARS-CoV-2, and increased risk of severe acute respiratory syndrome in COVID-19 patients, are present in the input datasets (14–16). To characterise the mutations in these susceptibility genes further, distinct mutation hotspots in the cohort can be visualised alongside the resulting amino acid (Figure 1B).

**Figure 1.**
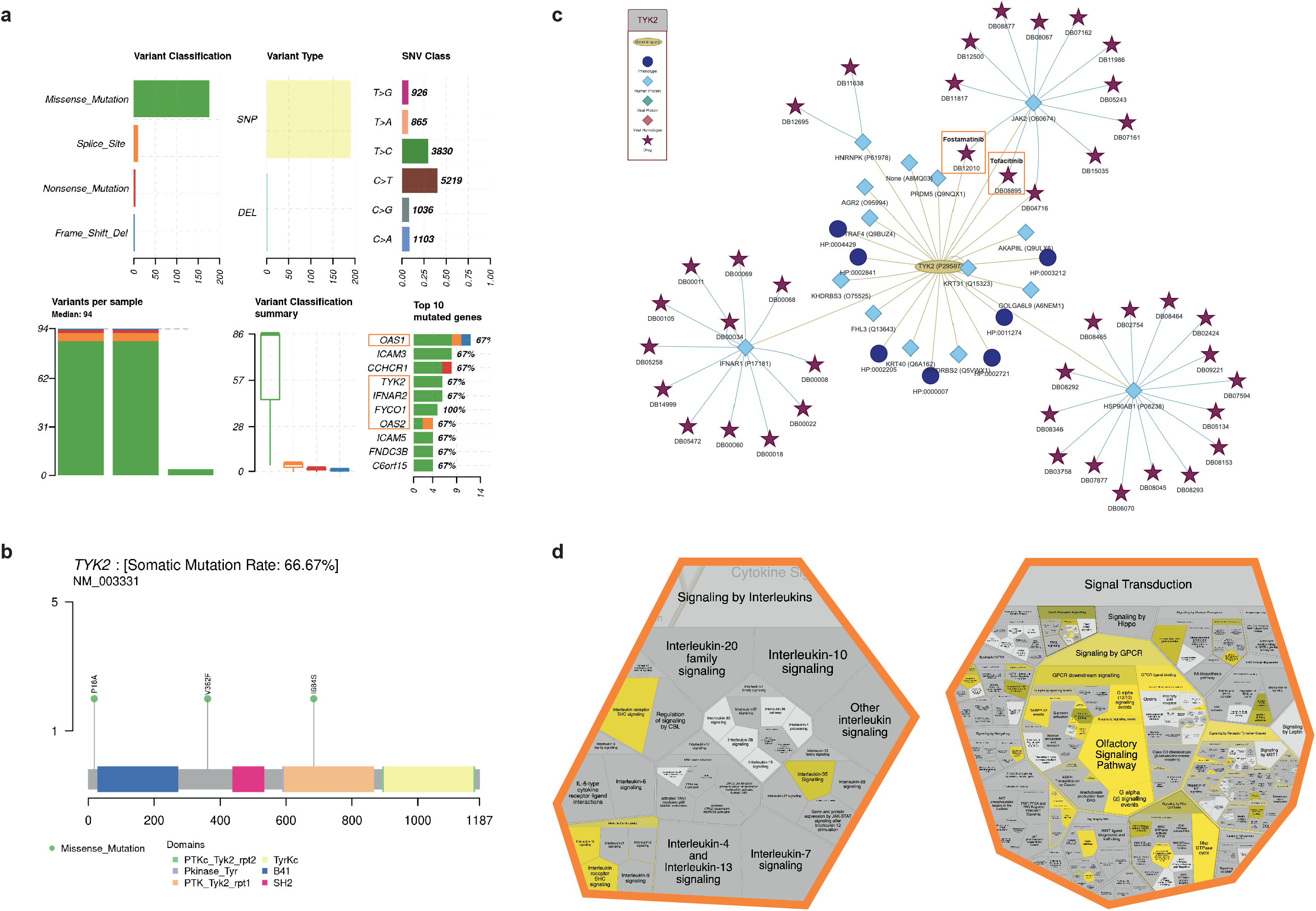
Analysis of sequence variations from The COVID-19 Host Genetics Initiative cohort. **(a)** A summarised view of the variant features in each dataset is provided, highlighting the variant classification distributions, SNV class and the most mutated genes across the cohorts. **(b)** The lollipop plot enables visualisation of mutation hotspots in a gene of interest with the amino acid changes for each gene labelled. **(c)** The protein-drug interaction network displays TYK2 with its associated phenotype, interacting proteins (targets) and drugs that target all proteins present. **(d)** The Voronoi diagram depicts enrichments in Reactome pathways. TYK2-associated pathways, such as *signaling by interleukins* and *signal transduction*, are reported as significantly enriched (p<0.05) in the input dataset.

SNPnexus COVID integrates data from IntAct Molecular Interaction Database, the Human Phenotype Ontology and DrugBank to generate networks that allow users to view proteins of interest, their associated phenotypes and drugs that target them.

Increased concentration of serum proinflammatory cytokines and chemokines has been correlated with adverse clinical outcomes in COVID-19 patients. Janus kinases (JAKs), such as TYK2, mediate intracellular signaling pathways employed by cytokines. As such, targeting the JAK-STAT signaling pathway with JAK inhibitors is being investigated for its ability to calm the cytokine release syndrome in COVID-19 patients (17).

Focusing on TYK2, SNPnexus COVID immediately identified TYK2 as a target for fostamatinib and tofacitinib (Figure 1C). Both of these drugs are being tested in clinical trials on hospitalised patients with COVID-19: fostamatinib to accelerate recovery and tofacitinib to suppress the pro-inflammatory response in patients with COVID-19 pneumonia (https://clinicaltrials.gov/ct2/show/NCT04579393, https://clinicaltrials.gov/ct2/show/NCT04469114).

The literature identifies anosmia and vascular promotion in COVID-19 patients (18,19). SNPnexus COVID highlights significant disruptions to GPCR signaling, affecting olfactory signaling pathways (Figure 1D).

SNPnexus COVID provides researchers with an intuitive web-based resource from which to explore their results, prioritise functionally and clinically relevant variants, and identify promising candidate drugs for repurposing.

## DISCUSSION

To maximise the usefulness of sequencing data in COVID-19 research, it is vital to have computational strategies in place capable of interpreting the data generated and ensuring that the clinically-relevant findings are translated back to the bedside. SNPnexus COVID provides researchers with access to extensive computational resources and facilitates the identification of novel biomarker and therapeutic targets in a timely manner. This optimises national efforts directed towards informing strategies for risk management within vulnerable populations and shaping clinical decision-making on candidate therapeutics.

SNPnexus COVID aims to remain abreast of evolving COVID research; the gene lists from GENCODE and Genomics England are continually updated within the tool. Furthermore, we welcome suggestions from our user community to expand and improve on the functionalities provided in this COVID-19 release.

## AVAILABILITY

SNPnexus COVID is freely available at https://www.snp-nexus.org/v4/covid/

